# SIngle cell level Genotyping Using scRna Data (SIGURD)

**DOI:** 10.1101/2024.07.16.603737

**Authors:** Martin Graßhoff, Milena Kalmer, Nicolas Chatain, Kim Kricheldorf, Angela Maurer, Ralf Weiskirchen, Steffen Koschmieder, Ivan G. Costa

**Affiliations:** Institute for Computational Genomics, RWTH Aachen University, Pauwelsstr. 30, 52074 Aachen, NRW, Germany; Department of Hematology, Oncology, Hemostaseology and Stem Cell Transplantation, RWTH Aachen University, Pauwelstr 30, 52074 Aachen, NRW, Germany; Center for Integrated Oncology Aachen Bonn Cologne Düsseldorf (CIO ABCD); Institute of Molecular Pathobiochemistry, Experimental Gene Therapy and Clinical Chemistry (IFMPEGKC), RWTH Aachen University, Pauwelstr 30, 52074 Aachen, NRW, Germany

## Abstract

**Motivation:** By accounting for variants within measured transcripts, it is possible to evaluate the status of somatic variants using single-cell RNA-sequencing (scRNA-seq) and to characterize their clonality. However, the sparsity (very few reads per transcript) or bias in protocols (favoring 3’ ends of the transcripts) makes the chance of capturing somatic variants very unlikely. This can be overcome by targeted sequencing or the use of mitochondrial variants as natural barcodes for clone identification. Currently, available computational tools focus on genotyping, but do not provide functionality for combined analysis of somatic and mitochondrial variants and functional analysis such as characterization of gene expression changes in detected clones.

**Results:** Here, we propose SIGURD, which is an R-based pipeline for the clonal analysis of single-cell RNA-seq data. This allows the quantification of clones by leveraging both somatic and mitochondrial variants. SIGURD also allows for functional analysis after clonal detection: association of clones with cell populations, detection of differentially expressed genes across clones and association of somatic and mitochondrial variants. Here, we demonstrate the power of SIGURD by analyzing single-cell data of colony-forming cells derived from patients with myeloproliferative neoplasms.

**Availability:** Code and tutorial of SIGURD are available at GitHub https://github.com/CostaLab/sigurd.

**Supplementary Information:** Supplementary data are available online.

## Introduction

The development of single-cell RNA-sequencing (scRNA-seq) methods allows the characterization of transcriptional changes in yet poorly characterized cell types^1,2^. This has been crucial to uncover cellular mechanisms related to diseases^3–6^. Clonal selection, i.e. the expansion of particular cell populations with malignant mutations, is a crucial mechanism in tumor cell development^7^. For example, in myeloproliferative neoplasms (MPNs), mutations in *JAK2, CALR* or *MPL*^7–10^ lead to the expansion of leukocytes, erythroid cells and/or megakaryocytes in the bone marrow. The measurement of the genotype of cells within an scRNA-seq experiment allowed the investigation of transcriptional changes in mutated versus non-mutated cells and provided important insights into the molecular mechanisms of cancer.

Theoretically, the mutation status can be measured using standard scRNA-seq protocols by detecting mutated reads covering the mutation locus. However, the sparsity and bias of standard scRNA-seq renders the chance of detecting mutations in coding regions extremely low, particularly with current protocols focusing on capturing the 3’ end and recovery of a fraction of transcripts per cell. To overcome this problem, modified scRNA-seq assays combined with advanced computational methods have been developed^11–14^. This allowed the categorization of cells by both transcriptional and mutational status. In turn, this enabled the detection of expanded cell type populations and the differential gene expression analysis between mutated and wild-type (WT) cells within a patient. For example, the amplification of the *CALR* locus in MPN samples increased the detection of mutated cells from 1.4% to 88.7% and indicated the association of *CALR* mutated cells with unfolded protein response and NKFB signaling when compared to WT^11^. Later, a similar approach was used to characterize *JAK2*V617F mutated cells^13^ detected an enrichment in erythroid progenitors. Both studies focused on experimental protocols, and relied on the use of custom/simple bioinformatics scripts for genotyping.

Recently, two computational genotyping methods (cellSNP-lite^15^ and VarTrix^16^) have been developed. These use read pile-ups obtained with short-read aligners to determine the number of mutated reads per position and cell. cellSNP-lite focuses on a time-efficient implementation for the detection of mutants in arbitrary regions. cellSNP-lite can only genotype single nucleotide polymorphisms (SNPs), and it only considers the first sequence of a unique molecular identifier (UMI) for genotyping to increase speed. VarTrix is a more general tool that also allows the detection of short indels. For genotyping, VarTrix performs a consensus variant call by considering all reads with the same UMI, which makes it more precise than cellSNP-lite. Due to the high computing time VarTrix can, in practice, be used only to genotype pre-selected genomic variants. Despite the power of these methods in the detection of variants in singlecells, they do not provide any support for quality checks or additional down-stream analysis of transcriptional data, such as the identification of clones and differential expression analysis of mutated vs. WT cells.

An alternative approach for single-cell genotyping is to explore mitochondrial (mt) variants (mtVars), serving as natural cellular barcodes for clones^12^. The fact that mtRNA transcripts are highly expressed makes the detection of mtVars easier than somatic variants (sVar). Moreover, amplification of mtRNAs in conjunction with standard 3’ scRNA-seq protocols and computational analysis improved the detection rate of clones by up to 50-fold^14^. However, the analysis of mtVars is not trivial, because they are non-functional. This can be solved by joint analysis of sVar and gene expression. For example, Caleb and colleagues demonstrated that characterized clones were both biased towards myeloid lineages and related to *TET2* variants in an individual with clonal hematopoiesis^14^. The Mitochondrial Alteration Enrichment and Genome Analysis Toolkit (MAEGATK)^14^ is a computational tool for genotyping mtVars from scRNA-seq. This tool uses aligned reads to detect all possible mtVars and uses a feature selection approach to filter mtVars that are specific to only a particular set of cells. As with previous genotyping tools, MAEGATK has its own output standards, and does not provide any integration with single-cell expression frameworks, such as Seurat^17^.

Altogether, computational analysis of genotyping data focuses on specific protocols (mitochondrial^14^ or somatic^13^ variants). Currently, we lack a framework for integrating mtVar and sVar with gene expression data.

### Our Approach

Motivated by this, we developed SIGURD (**SI**ngle cell level **G**enotyping **U**sing sc**R**na **D**ata), an R package designed to combine the genotyping information from both sVar and mtVar analysis from distinct genotyping tools and integrative analysis across distinct samples. SIGURD provides a pipeline with all necessary steps for the analysis of genotyping data, including: 1) candidate variant acquisition, 2) pre-processing and quality analysis of scRNA-seq with or without amplicon-based enriched sequencing, 3) cell-level genotyping via integration of several tools, and 4) representation of genotyping data in conjunction with the RNA expression data (**Fig. 1**). The last step allowed a seamless integration with Seurat for further down stream analyses. These include contrasting single-cell genotyping and transcriptional status, for example enrichment of mutations in cell types and cell type-specific differential expression analysis (WT vs. mutated cells). The genotyping information is saved as an R *SummarizedExperiment*^18^ object and can be added to a *Seurat*^19^ object as additional meta data. SIGURD can also consider candidate variants retrieved from data-bases such as the **C**atalogue **O**f **S**omatic **M**utations **I**n **C**ancer (COSMIC)^20^. This allowed us to focus on somatic candidate variants related to a particular disease entity, such as MPNs, which enhances both statistical power and lower computational requirements.

**Figure 1.**
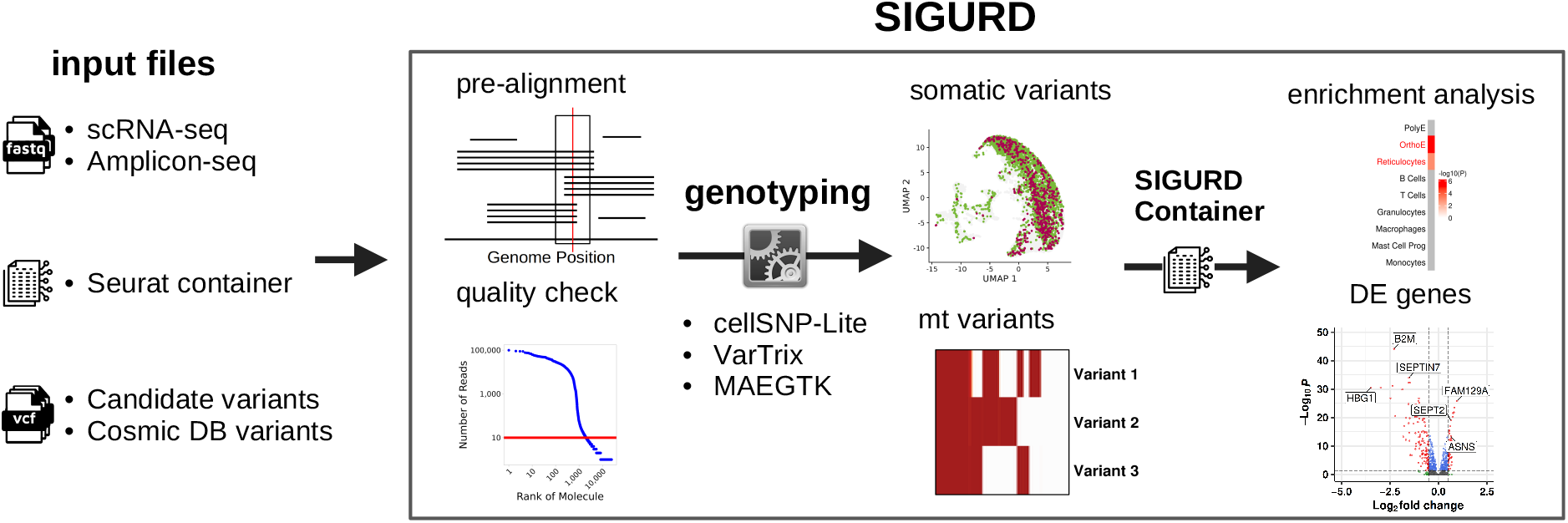
Overview of SIGURD. SIGURD receives as input fastq files with reads generated in an amplicon or single-cell sequencing experiment, as well as a Seurat container with pre-processed scRNA-seq data. Additional inputs are VCF files with variants. Then, SIGURD performs a pre-alignment and quality control to detect valid molecules. Subsequently, it allows the use of distinct single-cell genotyping tools (VarTrix, cellSNP-lite and MAEGATK) and supports the detection of clonal cells. Results are integrated in a SIGURD data container and they can be analyzed (together with the scRNA-seq container) for tasks such as differential expression or enrichment analysis between clones.

To demonstrate the power of SIGURD, we present here a case study on the analysis of scRNA-seq in clonogenic cells from six MPN patients with polycythemia vera (PV), post-PV-myelofibrosis, and essential thrombocythemia (ET)^21^, and three healthy controls (HC)^22^. We also amplified the *JAK2*V617F locus, as previously described^13,23^, the driver mutation in all analyzed MPN samples. We used this comprehensive data, which includes novel ET samples not analyzed previously^22^, to show how SIGURD can combine mtVar and sVar analyses to characterize clonally associated transcriptional changes.

## Methods

The SIGURD pipeline receives as input a set of BAM files related to the single-cell and amplicon sequencing. Optionally, the user can provide a set of candidate regions for genotyping. Next, a pre-alignment and quality check analysis was performed to consider the coverage of reads in the variants and to support the selection of parameters associated with valid transcripts. Subsequently, it allows the use of a combination of genotyping tools (VarTrix, cellSNP-lite and MAEGATK). Finally, it combines the results of genotyping tools by producing a *Seurat*-compatible container with both genotyping and single-cell data for further integrative analysis of the genotyping and transcriptome data.

### Input

The genotyping data analysis requires several input files. This includes BAM files from scRNA-seq and amplicon sequencing (for either or both sVar/mtVar) and an scRNA-seq container processed with the scRNA-seq pipeline using Seurat^24^.

Regarding sVars, cellSNP-lite or VarTrix can align and perform variant detection in the entire genome. Such genome-wide genotyping requires a long computing time and/or will find non-functional genotypes. To address this, SIGURD supports the selection of a list of candidate variants, obtained from the COSMIC database^20^. The COSMIC data-base contains information on millions of cancer-related variants. This generates a variant call format (VCF) file, which can be used as an input for SIGURD.

### Initial Alignment and Quality Check

Before genotyping, SIGURD performs a pre-alignment and a quality control (QC). The combination of a cell barcode and UMI can be referred to as a molecule, i.e. a unique transcript measured in a specific cell. To avoid sequencing artifacts, genotyping tools should only consider molecules with sufficient reads that support a particular variant. This should be done in a sample- and protocol-specific manner, with the help of QC plots, as described below. In addition, only cells detected in the matching scRNA-seq (count matrix) were considered and reads with a mapping quality <30 were ignored.

SIGURD performs a pre-alignment with STAR^25^. It then generates QC plots to select parameters for the detection of valid molecules for somatic and mitochondrial variants. To define the cut-off number of reads per molecule, SIGURD provides plots (**Fig. 2A** and **Fig. S1**). A visual inspection of these plots indicates knees (drops in distributions) suggesting optimal threshold values. Importantly, this cut-off has an impact on the number of detected genotyped cells (**Fig. 2B**).

**Figure 2.**
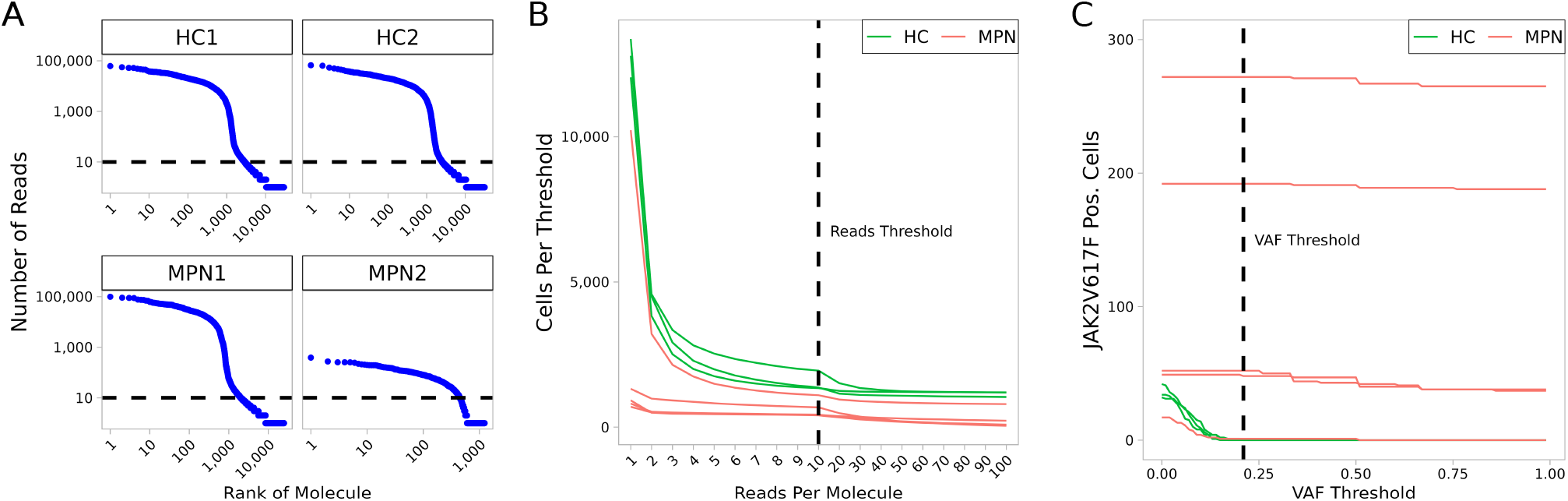
Quality Check Plots. **A** Plot for number of reads per molecule (UMI) (y-axis) vs. the rank of the molecule (x-axis) for *JAK2*V617F amplicon sequencing of selected MPN and HC samples. A knee in this plot can indicate a threshold for considering valid molecules in a particular library, as indicated with the dashed line. **B** The number of cells for various threshold of reads per molecule for selected libraries. The dashed line indicates the threshold. **C** The number of cells for various thresholds of variant allele frequency (VAF). The dashed line indicates a (VAF) threshold (21%) below which positive cells are not detected in HC samples.

An alternative approach to increase cell recovery is to select a more lenient threshold of reads per molecule, but filter cells by the variant allele frequency (VAF) in the case that disease and healthy control samples are available. One can choose a VAF with no positive cells (false positives) in healthy controls (**Fig. 2C**). Another important plot is the visualization of the read coverage at a particular locus (**Fig. S2**). This illustrates if and how efficiently the amplification protocol worked.

The pre-processing will result in a filtered BAM file, that only contains high-quality reads confidently mapped to annotated cells.

### Genotyping

The genotyping module receives the pre-selected variants and filtered aligned reads as inputs. The genotyping can be performed using any combination of the following tools: VarTrix, cellSNP-lite and MAEGATK. For genotyping specific sVars, the aforementioned VCF file with candidate variants is required.

**VarTrix**^16^ evaluates aligned reads from a BAM file for each variant provided as input. For this, a VCF file containing the genomic information of the variants is required as input. The reads are locally re-aligned using the Smith-Waterman algorithm^16^. There are two output modes: consensus and alternative fraction. In the consensus mode, for each cell and variant, the output is either 0 when no reads cover the variant (NoCall), 1 when only reads supporting the reference sequence cover the variant (Ref), 2 when only reads supporting the alternative sequence cover the variant (Alt), or 3 when a mixture of alternative and reference reads covers the variant (Both). In alternative fraction mode, VarTrix produces a matrix that indicates the percentage of alternative reads per cell and variant. In combination with the previous options, VarTrix can be run in UMI mode (*–umi*). In UMI mode, VarTrix takes advantage of the sequencing depth of amplicon assays by taking the consensus of all reads per molecule, that is it only considers molecules with more than 90 % of the reads supporting a sequence. We strongly recommend the use of the UMI mode.

**cellSNP-lite**^15^ was designed to genotype cells at a high speed and low memory usage. This allows genotyping of full genomes, but only SNPs can be detected. A simple genotyping strategy, for example only the sequence of the first read is used, making cellSNP-lite very efficient, but potentially error-prone. The output from cellSNP-lite is a VCF file. In addition to standard VCF fields, the files contains a column for each cell with its genotype, read depth and the number reads supporting each possible nucleotide. When analyzing a public data set^26^, cellSNP-lite was six times faster than VarTrix, while using only 10% of the memory^15^, that is cellSNP-lite required only 45 minutes and less than 1 GB memory in the analysis of standard scRNA-seq data^15^. However, by not considering the consensus sequences in reads with the same UMI, cellSNP-lite ignores the great depth of amplicon sequencing. In such cases, VarTrix is preferable.

**MAEGATK**^14^ was designed to analyze clones by exploring the mitochondrial variants. MAEGATK is an extension of the Mitochondrial Genome Analysis Toolkit (MGATK)^27^, designed for the analysis of scATAC-seq data. MAEGATK requires, as input, a BAM file and a list of cell barcodes. Reads are then grouped into molecules by UMI, cell barcodes and the aligned positions. The reads are then collapsed, i.e. a consensus sequence for each group of reads is formed by identifying the most likely nucleotide at each position by majority vote. By default, SIGURD only considers molecules with at least three reads^14^.

Clonality-related mitochondrial variants of interest (mtVOIs) must be detected de-novo and in an individual-specific manner^14,28^. To facilitate this, SIGURD implements two mtVOI selection approaches^14^.

In the unsupervised approach, SIGURD considers variants with a minimum VAF of *X* (default 90%) in the top quantile (default 90%) and a maximum VAF of *Y* (default 10%) in the bottom quantile (default 10%). This procedure guarantees that the candidate variant is present in at least a significant percentage but not in all cells in the sample, as exemplified in **Fig. 3**. In the supervised approach, variants are selected by considering cell information, for example time-point, treatment conditions or scRNA-seq derived cell types when measured over a given donor/patient. For this, SIGURD first finds initial variants by using the unsupervised procedure in only one of the conditions. Next, it selects variants with an average VAF *x* times higher (default=5) in the condition of interest vs. the other cells. After mtVOIs are selected, SIGURD allows their visualization as a heat-map. Moreover, SIGURD allows the definition of clones by considering all combinations of mtVOIs. By considering the fraction of cells in a clone, clonal expansion can be detected. For further down stream analysis, SIGURD performs a filter by considering frequent clones (**Fig. 6C**).

**Figure 3.**
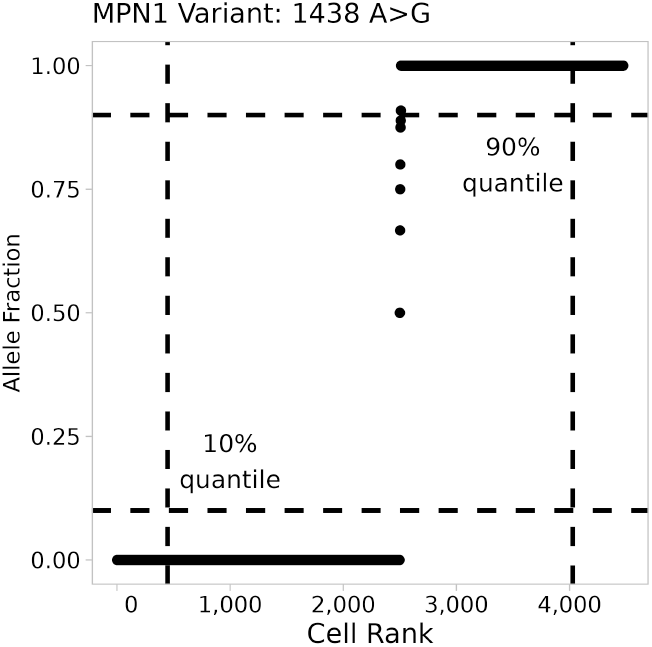
MT Variant Analysis. Plot displaying the VAF of cells for a given mtVar (y-axis) sorted by increasing VAF (x-axis). The name of the mtVar (1438 A*>*G) indicates the position in the mitochondrial genome and the detected base substitution. Dashed lines indicates the quantile thresholds used by SIGURD.

### Integrative Analysis of Expression and Genotypes

A typical experimental design will have scRNA-seq and amplicon data for several individuals/patients over distinct conditions: treatment or non-treatment, or time series design. Therefore, variant information needs to be combined per individual. To facilitate this, SIGURD requires an experimental design matrix, in which users define individuals and conditions as their respective input files (see **Table S1** for an example).

For every individual, SIGURD reads the genotyping results and stores them in a *SummarizedExperiment* R object. This object holds the information if a variant was detected (Alt.), non-detected (Ref.) or if no call was possible (No Call). For every entry, it also records the information of molecules supporting the calls, the fraction of the mutated molecules, and read coverage. For the MAEGATK results, the object also contains the positional sequencing quality and the strand concordance. Additional meta data, such as cell types derived from scRNA-seq analysis, can be added.

Next, SIGURD allows the use of filters to improve genotyping calls. The first filter includes a minimum VAF value, because a low VAF is possibly related to artifacts/false positives. Cells that do not pass this threshold, which is set by default as 0, are defined as NoCall (**Fig. 2C**). Finally, SIGURD adds the final genotype information to a *Seurat* object to relate the clones with gene expression and cell type (from the scRNA-seq object) to SIGURD object.

With the combined information, the user can perform several analyses of the data. For sVars, a functionality of SIGURD is the estimation of the VAF per sample, which can be compared with the mutant VAF determined via DNA-sequencing. The same information can be compared for each cell type to verify whether the VAF is cell type-specific. The genotyping information can be used as a cell annotation variable, which can be utilized for visualization on a UMAP. Another functionality is the use enrichment tests (based on Fisher’s Exact Test) to check if an sVar is associated with a cell type or a condition (treatment vs. non-treatment) (**Fig. 4**).

**Figure 4.**
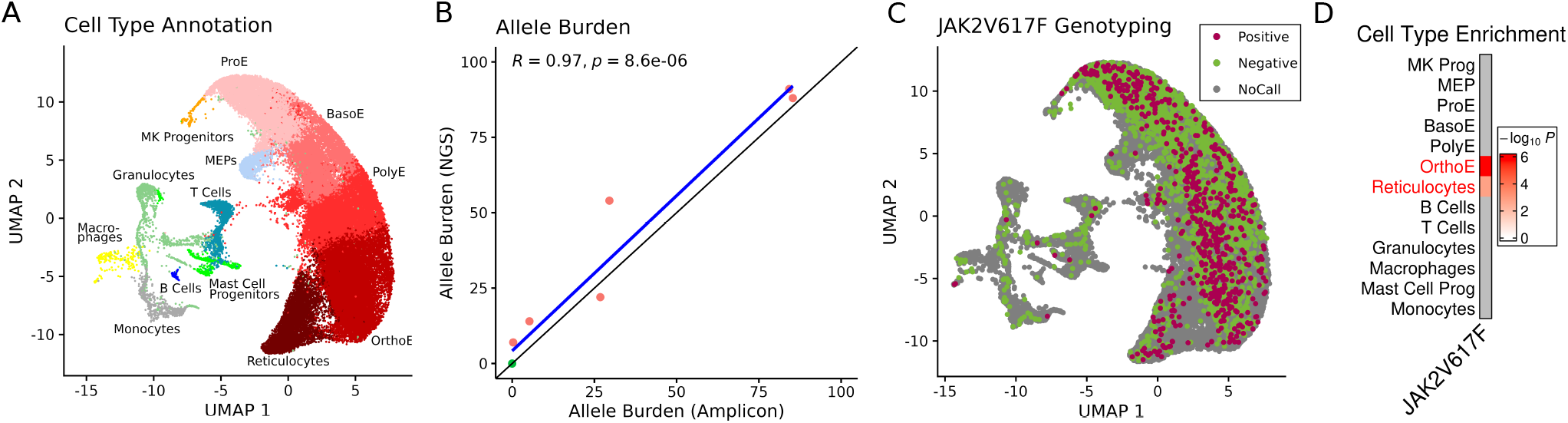
*JAK2*V617F Genotyping. **A** UMAP with cell types detected in single-cell experiments. **B** Scatter plot of the *JAK2*V617F allele burden measured by amplicon data (x-axis) and DNA-sequencing (y-axis). We correlated the values using Pearson Correlation Coefficient. **C** UMAP displaying *JAK2*V617F positive, negative and no-called cells. **D** Cell type enrichment for the *JAK2*V617F positive (vs. negative) cells by cell types (adjusted P-values; Fisher’s Exact test).

Finally, differential expression analysis (DE) is performed, e.g. comparing WT (Ref.) and Mutated (Alt.) cells for a given cluster using standard DE analysis and Gene Set Enrichment Analysis (GSEA) methods **Fig. 5**). DE analysis used the Wilcoxon Rank Sum Test^29^ and *p*-values were adjusted using the Bonferroni method. GSEA was based on a univariate linear model using decoupler^30^ and *p*-values were adjusted using the Benjamini-Hochberg correction^31^.

**Figure 5.**
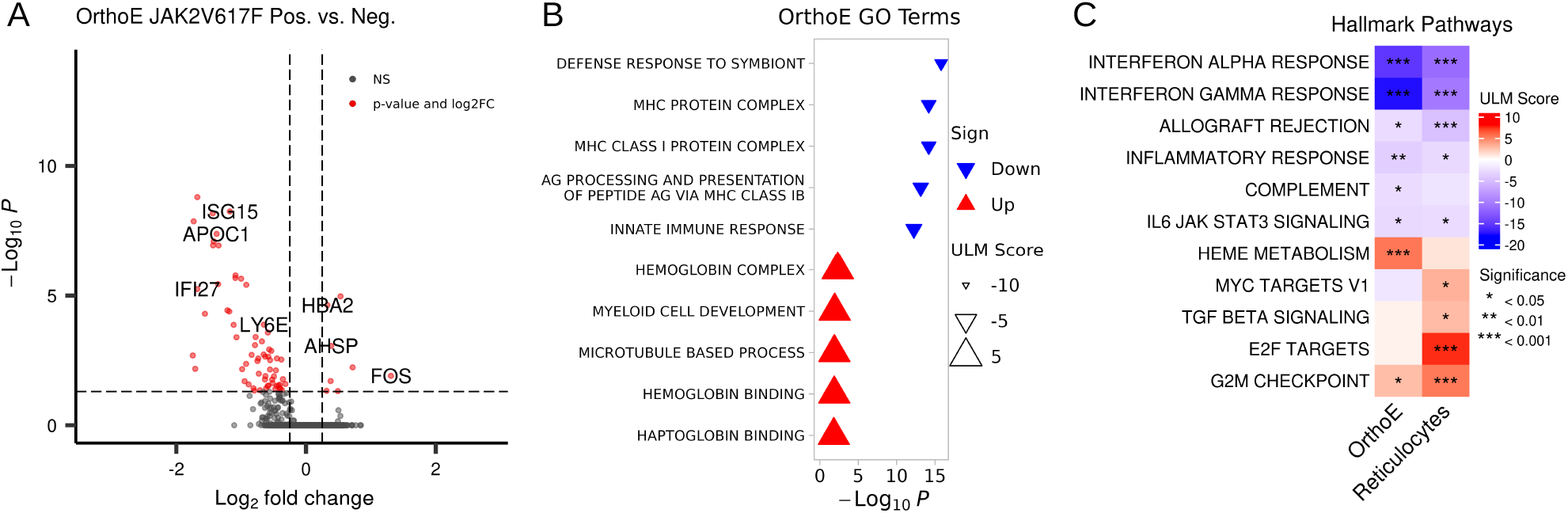
Differential Gene Expression Analysis. **A** Volcano plot showing the DEGs for the comparison *JAK2*V617F vs. WT for the OrthoE cells. **B** ULM results for GO terms (top 5 up and down) for OrthoE DE analysis. **C** ULM for hallmark gene sets for OrthoE and reticulocytes. Only pathways with at least one enrichment (adjusted P-value <0.05) are displayed.

Finally, SIGURD uses the combination of mtVOIs to define clones. We can then explore ecological measures to measure clone diversity, as low diversity is indicative of clonality. SIGURD allows the use of classical measures such as Shannon Entropy^32^, i.e.

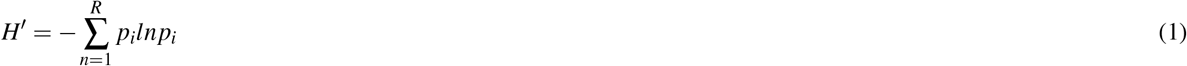

where *p*_*i*_ is the probability of a cell being part of clone *i*. We can also define the effective species^33^ values as:

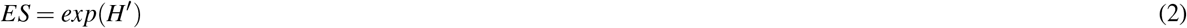

### 0.1 Transcriptional and Cellular Changes of JAK2V617F Positive Cells Material and Methods

#### Cell Cultivation and Treatment

Treatment and analysis of the cells were performed as described previously in Kalmer et al^22^. Briefly, peripheral blood mononuclear cells were isolated and seeded in a colony-forming unit assay. Cells were harvested using pre-warmed PBS and applied to scRNA-seq following Chromium Next GEM Single Cell 3’ Reagent Kits v3.1 User Guide Rev D. Sequencing was done on a NextSeq500 using NextSeq500/550 Mid Output Kit v2.5 (150 cycles) or a NextSeq500/550 High Output Kit v2.5 (150 cycles). Locus specific amplification of *JAK2*V617F was performed as following Liu et al^23^, labeled using Sample Index Kit T Set A and sequenced on a NextSeq500/550 Mid Output Kit v2.5 (150 cycles) with 50% PhiX. MPN1, MPN2, MPN3, WT1, WT2 and WT3 are the untreated samples as described our previous paper^22^. The samples MPN4, MPN5, and MPN6 are new samples that have been processed together with the original data set. MPN1, MPN2, and MPN3 are PV samples, MPN4 was diagnosed with post-PV-myelofibrosis and MPN5 and MPN6 suffer from ET. None of the patients received cytoreductive treatment at the time of sampling. We have scRNA-seq data for all samples and *JAK2*V617F amplicon sequencing for all samples but for MPN6.

#### scRNAseq Processing

We pre-processed the scRNA-seq data for each library with a cellranger pipeline (version 3.1.0) using the default parameters (genome version GRCh38). Data integration and clustering were performed using Seurat (version 4.2.0). We removed cells with 400 or fewer expressed genes, more than 40,000 reads in total, and genes expressed in less than 5 cells. Doublet detection and removal were performed with DoubletFinder^34^. Following data integration and principal component (PC) analysis, we constructed a shared nearest neighbor graph using the first 30 PCs and a clustering resolution of 0.5. Cell cycle scoring was performed using the *CellCycleScoring* function from Seurat.

Next, we ran SIGURD for the scRNA-seq and amplicon data. The cells were genotyped for sVars using VarTrix in UMI mode with a minimum mapping quality of 30. To increase cell recovery, we used a lenient threshold of 1 read per molecule. We only considered cells with a VAF >20%. MAEGATK was used for mtVars, with a minimum of 3 barcode reads and a minimum alignment quality of 30. We retrieved differentially expressed genes using Seurat’s *FindMarkers* function with the Wilcoxon Rank Sum Test. Only genes expressed in at least 10% of the cells in one of the groups were tested. *P*-values were adjusted using the Bonferroni correction.

## Results

Here, we describe the application of SIGURD in single-cell data of clonogenic cells derived from MPN patients and healthy donors. First, we describe the genotyping results of amplicon sequencing, amplifying the *JAK2*V617F mutation in these samples. Next, we show results of clonal analysis by exploring mtVars. Finally, we present the combination of these distinct genotyping strategies.

### Genotyping of JAK2V617F Positive Cells

Single-cell experiments of three healthy controls and six MPN samples captured 57,905 high-quality cells (average of 1,297 genes per cell)^19^. We next used the available single-cell data of blood cells^1,35,36^ to annotate the cells. This resulted in a large cluster, which contained erythroid and megakaryocytic cells (megakaryocyte progenitors (MK Prog), megakaryocyte-erythroid progenitors (MEP), pro-erythroblasts (ProE), basophilic (BasoE), polychromatic (PolyE), orthochromatic (OrthoE) erythroblasts, and reticulocytes; **Fig. 4A**). In addition, smaller lymphoid (T and B cells) and myeloid (granulocytes, macrophages, mast cell progenitors, and monocytes) cell populations were also recovered. See **Fig. S3** for marker gene expression in our data-set. Next, we performed *JAK2*V617F amplicon based genotyping for five of six MPN samples and all three healthy controls. We used VarTrix (as opposed to cellSNP-lite) because it considers all reads of a molecule for genotyping, and only *JAK2*V617F needed genotyping. As a first step, we performed QC analysis of the amplicon sequencing to check the amplification of *JAK2* transcripts and cut-offs to delineate *JAK2*V617F positive cells. As shown in **Fig. S2**, amplicon sequencing increased read recovery 8-fold in most libraries.

Our initial quality check analysis indicated that 52,067 cells (out of 57,905) had at least one molecule covering the *JAK2*V617F locus (**Fig. 2B**), but the number of cells dropped quickly when more reads per molecule were required. Therefore, we used a lenient threshold of one read per molecule. Next, we checked the number of detected positive cells in all libraries for distinct VAF thresholds (**Fig. 2C**). We selected the minimum threshold of 21%, as this threshold did not detect any false positives (*JAK2*V617F positives) in the healthy control samples. As an additional check, we correlated the VAF detected using the scRNA data and NGS-sequencing (**Fig. 4B**). This indicated a high correlation (0.97; n=8), which supports the high agreement with the scRNA-seq based genotyping with DNA-sequencing. Altogether, SIGURD detected 603 *JAK2*V617F positive and 2,616 negative cells.

By combining expression and genotyping data, SIGURD allows to detect cellular and gene expression changes related to mutations. UMAP indicated that *JAK2*V617F positive cells were particularly abundant among erythroid cells (**Fig. 4C**). A more principled way to analyze this is by using an enrichment test (Fisher’s Exact test), which was performed and further indicated that *JAK2*V617F cells were particularly enriched in mature erythroid fractions (OrthoE and reticulocytes; **Fig. 4D**). To determine the impact of the *JAK2*V617F mutation on the gene expression of cells, we contrasted the expression profiles of positive and negative cells. To retain statistical power, we focused on cell types with a minimum number of cells (*>*50) per condition (Supp. **Tab. S2**). Of particular interest were OrthoE and reticulocytes, as these were enriched with *JAK2*V617F positive cells.

DE analysis detected 8 up- and 64 down-regulated genes in OrthoE (**Fig. 5A**), whereas no DE genes were found in reticulocytes (**Fig. S4**). Down-regulated genes (in OrthoE) included genes related to MHC protein complex and interferon response genes, while up-regulated genes included hemoglobin-related genes such as HBA2. The absence of DE genes in reticulocytes can be explained by the lower cell recovery (74 genotyped cells vs. 345 for OrthoE). An alternative analysis to address this is GSEA, which considers fold changes of all measured genes. This revealed common pathways between OrthoE and reticulocytes, such as an increase in heme metabolism and G2M checkpoints as well as a down-regulation of interferon response genes.

### Genotyping with Mitochondrial Variants

Next, we explored mtVars to identify clones in these cells using scRNA-seq data. The QC analysis indicated good cell recovery (16,777 HC and 40,559 MPN cells) when 3 reads per cell were used (**Fig. S5**). We next detected mtVOIs using an unsupervised approach and standard parameters (as shown in **Fig. 3**). This approach resulted in the selection of 3-14 mtVOIs per sample (**Figs. 6** and **S6**). SIGURD adopts the definition that a combination of variants defines a clone, in contrast to previous reports^14^, which used mtVOIs only. Next, we only considered clones covering at least a fraction of cells (*>*5%). With this, we detected 2-9 expanded clones in MPN, and 0-1 expanded clones in HC (**Fig. S6**). Cells with no positive mtVOIs (white columns in **Fig. 6A-B**) were annotated “negatives” and were disregarded in the downstream analyses. In these cells, mtVar detection failed, possibly due to low read coverage. The higher clonal expansion in MPN samples compared to HC is illustrated in **Fig. 6A-C** and **Fig. S6**. Ecology measures^32,33,37^ provide a more formal way to evaluate the diversity of species (clones) in a population. Indeed, we observed a clear drop in diversity in the MPN samples compared to control (**Fig. 6D**). This confirmed the clonal expansion of disease lineages, as expected in clonal hematopoiesis, a feature of MPNs.

**Figure 6.**
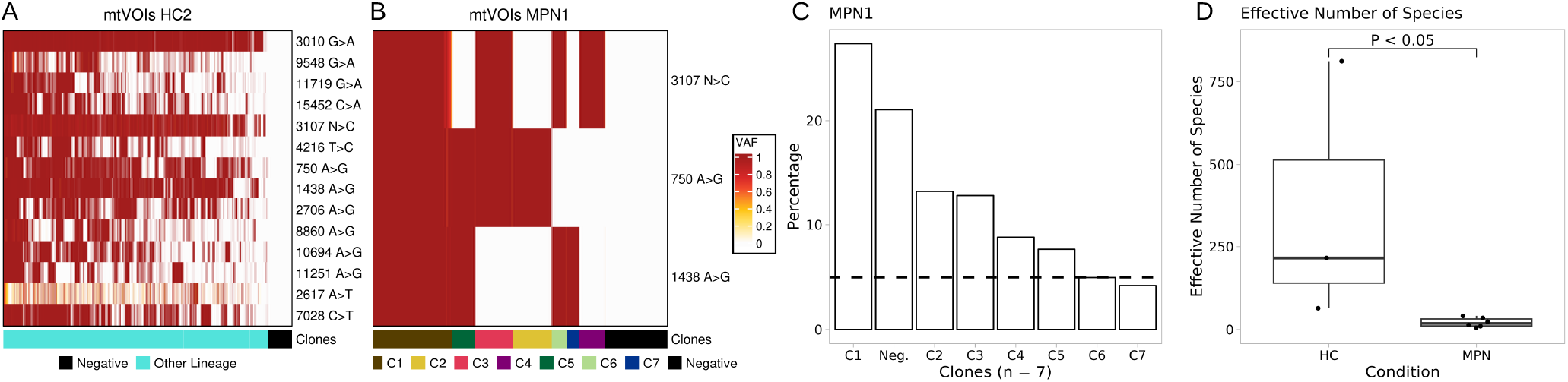
Mitochondrial Genotyping. **A-B** Heat maps with selected mtVOIs for samples HC2 and MPN1. The color gradient indicates the VAF. Moreover, combination of variants can be used to define clones, as indicated in the color bar below the heat map. “Negative” indicates cells with no detected mtVOI. **C** An important step in downstream analysis is to only consider clones covering at least a proportion of cells (5%, indicated by the dashed line). Each clone is a unique combination of mtVars, while the Negative clone is comprised of cells without any detected mtVar. **D** The effective number of species per condition. Comparison was performed using the Wilcoxon-Mann-Whitney test.

The functional characterization of clones derived from mtVars is not trivial, as we lack information on somatic mutations driving functional changes in these clones. In this case, we did not find any statistical association between the clones and *JAK2*V617F cells. However, the changes in cellular composition and gene expression can shed some light on this. We present in **Fig. S7** cell type enrichment and GSEA for the three largest clones (C1-C3) per MPN sample. For example, C1 cells in MPN1 were enriched in early erythrocyte cells (ProE, BasoE), while clones C2 and C3 were enriched in late erythroid cells (OrthoE, reticulocytes). When considering gene expression changes in these clones (GSEA), we observed higher expression of heme metabolism genes in C2-C3, but higher expression of MYC target genes in C1. Similar molecular profiles were found in MPN2, where C1 and C2 were mostly related to late erythroid cells and C3 to early erythroid cells.

To ensure that these differences were not only driven by cell composition, analysis of a clone should be performed by comparing cells per cell type. For MPN1 and C1, both PolyE and ProE showed significantly lower heme metabolism (**Fig. 7**). C1 PolyE also showed a reduction in interferon response genes, while ProE had an increased expression for G2M checkpoint, MYC targets, and oxidative phosphorylation pathways. In C2, OrthoE showed an increase in heme metabolism, interferon response and IL2 STAT5 signaling, while reticulocytes showed an increase in heme metabolism and a reduction in cell cycle pathways. In C3, we observed an increase in heme metabolism for PolyE and OrthoE. This diversity in differentially regulated pathways indicates that these molecular changes were driven by clonality.

**Figure 7.**
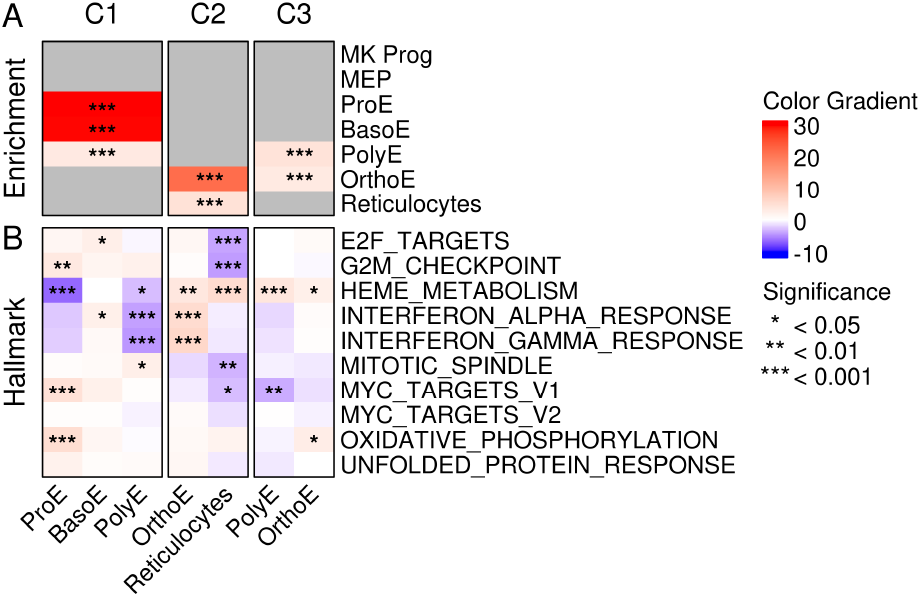
Mitochondrial Genotyping and Expression Changes. **A** Heat map with cell type enrichment per clone (*−*log_1_0 p-value) for the MPN1 sample. **B** GSEA (Hallmark pathways) values (ULM scores) on MPN1 clones of significantly enriched cell types from **A**). The same color scale was used for both results.

The procedure used for mtVar selection focused on variants identified in a large proportion of cells (*>*10%), as default by MAEGATK^14^. We reasoned that by using a more lenient threshold (*>*20 cells), we could identify mtVars related to *JAK2*V617F. Indeed, this procedure led to the identification of three mtVars in sample MPN2. Assuming that cells positive for these variants were *JAK2*V617F positive, this increased our cellular capture from 194 to 802 cells. GSEA analysis indicated similar molecular signatures in these cells, which supports similar molecular changes as in *JAK2*V617F cells (**Fig. S8**). These results further support the power of mtVars to improve genotyping of cells.

## Discussion

Here, we present SIGURD, a framework for analyzing single-cell genotyping data. SIGURD takes advantage of several genotyping methods for either somatic or mitochondrial variants, and allows the combined analysis of clonal/genotyping information with transcriptional information in the respective scRNA-seq data. This allowed us to gain insight into different types of variants, whereas previous approaches have focused on specific types of mutations, e.g. somatic variants^11^ or mitochondrial variants^14,27^. Our framework builds upon these previous and own publications^22^ to provide an end-to-end analysis of single-cell genotyping and gene expression. This includes the processes of initial data acquisition, filtering and quality control, detection of variants and integrative analysis.

We demonstrate the use of SIGURD in scRNA-seq and amplicon sequencing data of a driver mutation (*JAK2*V617F) in clonogenic cells from MPN patients. Combined analysis of *JAK2*V617F genotyping and gene expression indicated an enrichment of mutated cells in late erythroid cells and the up-regulation of heme metabolism genes. These findings validate each other, as more mature erythroid cell types show a higher expression of heme metabolism^38^.

Regarding mitochondrial variants, the comparative analysis indicated an expansion of clonal lineages within MPN patients compared to control. Interestingly, all MPN samples exhibited a mixture of clones enriched for later or early erythroid cells highlighting the heterogeneity of clones and the importance of their analysis. GSEA indicated that MYC target pathways were enriched in early erythroid cells, supporting expansion of these populations. Late erythroid cells have a similar signature as *JAK2*V617F positive cells, such as increase in heme metabolism. We did not observe any association of *JAK2*V617F positive cells and mitochondrially detected clones. An explanation for this discrepancy could be low cell recovery of mitochondrial variants in our data (52%). Note that previous work also used amplicon sequencing to enrich for mitochondrial reads^14,27^. However, when we used modified criteria to detect mitochondrial variants, we were able to identify variants associated to *JAK2*V617F cells in one sample. This supports the need for further studies on selection of mitochondrial variants of interest. Altogether, our analysis has shown how SIGURD can build upon genotyping and scRNA-seq, providing an in-depth analysis of multi-modal genotyping and transcriptomic data.

In the future, we will expand the described methods to include spatial data, such as spotted arrays^39^. This would allow the characterization of spatial organization of malignant clones. The sparsity of spatial data, which is greater than that scRNA-seq, requires the improvement of genotyping algorithms. Moreover, the fact that spotted arrays do not achieve cellular resolution requires novel methods for the selection of VOIs, which consider a mixture of cell populations.

## Data and Software Availability

We also deposited pre-processed Seurat and SIGURD objects in Zenodo (https://zenodo.org). The temporary link for reviewers is: https://zenodo.org/ Code and sample scripts of SIGURD are available at GitHub https://github.com/CostaLab/sigurd.

## Acknowledgments

This work was supported by the Genomics Facility of the Interdisciplinary Center for Clinical Research (IZKF) Aachen within the Faculty of Medicine at RWTH Aachen University. Biomaterial samples were provided by the RWTH centralized Biomaterial Bank Aachen (RWTH cBMB, Aachen, Germany) in accordance with the regulations of the Biomaterial Bank and the approval of the ethics committee of the medical faculty, RWTH Aachen. Part of this work was generated within the doctoral thesis works of MK and MG.

## Author contributions

MG, MK, SK and IC have designed the study, analyzed data and revised the manuscript. MG performed all programming tasks and wrote the first version of the manuscript. MK has performed all wet-lab work and sequencing experiments. All authors have revised and approved the final version of the manuscript.

