## Supplementary Information for "SIngle cell level Genotyping Using scRna Data (SIGURD)"

### 1 Supplementary Information

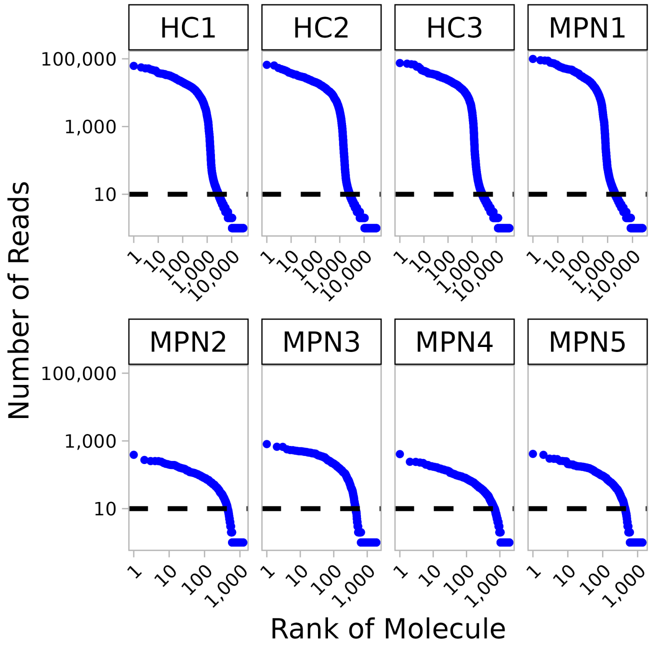


**Figure S1. Quality Check Plots** Plot for number of reads per molecule (UMI) (y-axis) vs. the rank of the molecule (x-axis) for *JAK2*V617F amplicon sequencing. A knee in this plot can indicate a threshold for considering valid molecules in a particular library, as indicated with the dashed line.

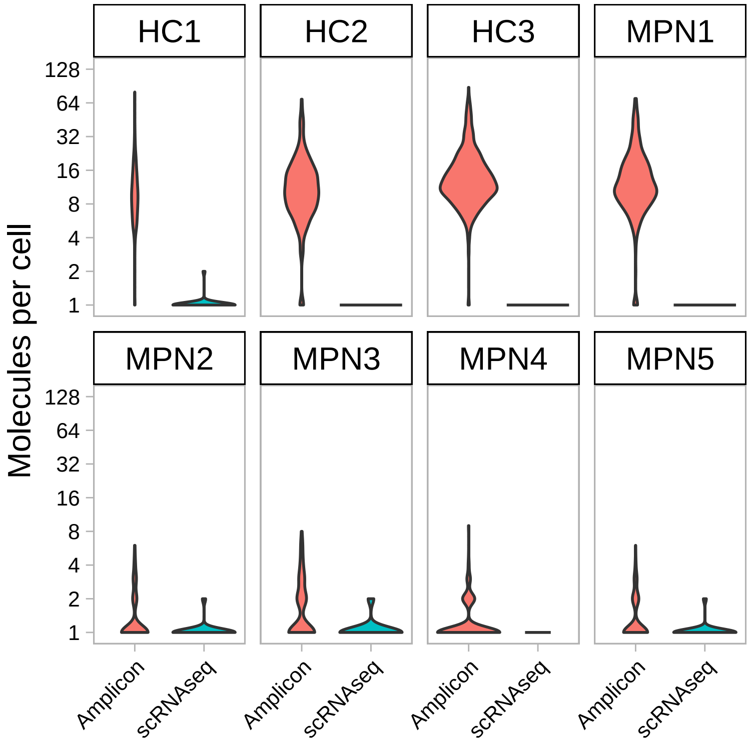


**Figure S2.** **Quality Check Plots *JAK2V617F* & Genotyping.** The number of molecules per cell displayed as a violin plot. The number of molecules per cell for the standard 10X scRNAseq data is very low and only one molecule per cell is detected. In these cases, the violin appears as a line.

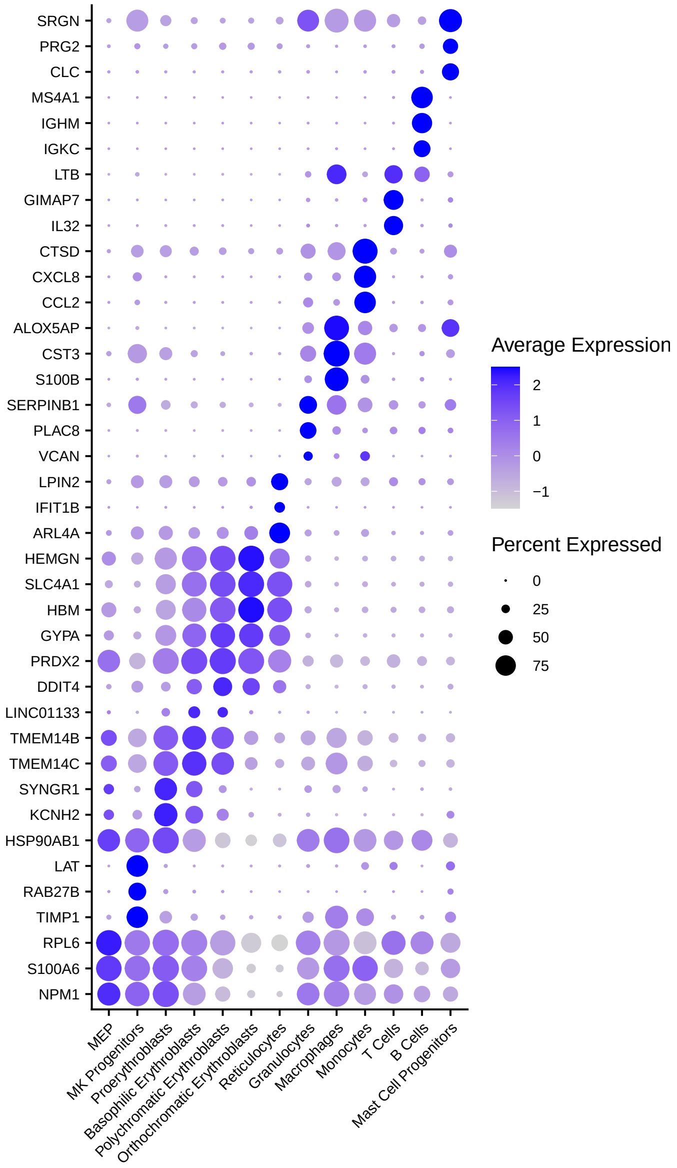


**Figure S3.** **Marker Genes per cell type.** The dot plot shows the top 3 marker genes used for the annotation. This expression pattern was used in conjunction with the Erythron Data Base[1] to conduct the cell type annotation.

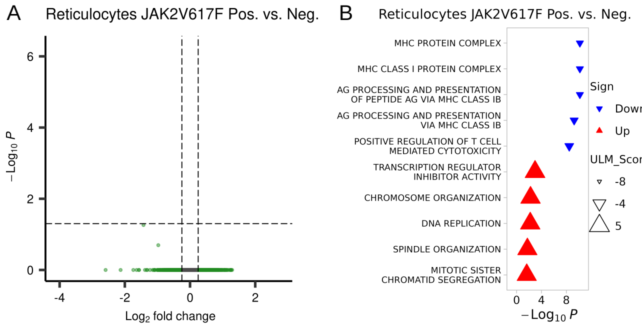


**Figure S4.** **DEGs reticulocytes *JAK2V617F* Pos. vs. Neg.** **A** A scatter plot showing the DEGs for the comparison *JAK2*V617F Pos. vs. Neg. for reticulocytes. **B** Top 5 up and down regulated GO terms for the DEGs.

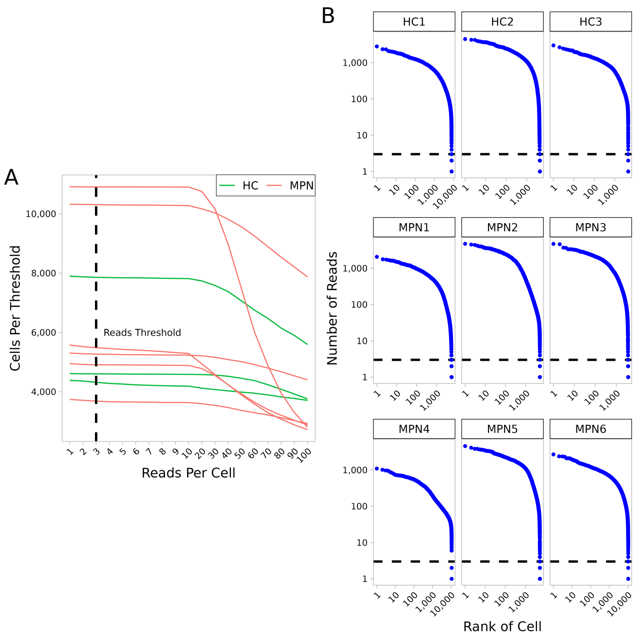


**Figure S5.** **Quality of MT sequencing. A** The number of cells retained for a given reads per molecule threshold. Cells without any molecules would not be analyzed in the MT analysis. **B** Number of reads per molecule per sample.

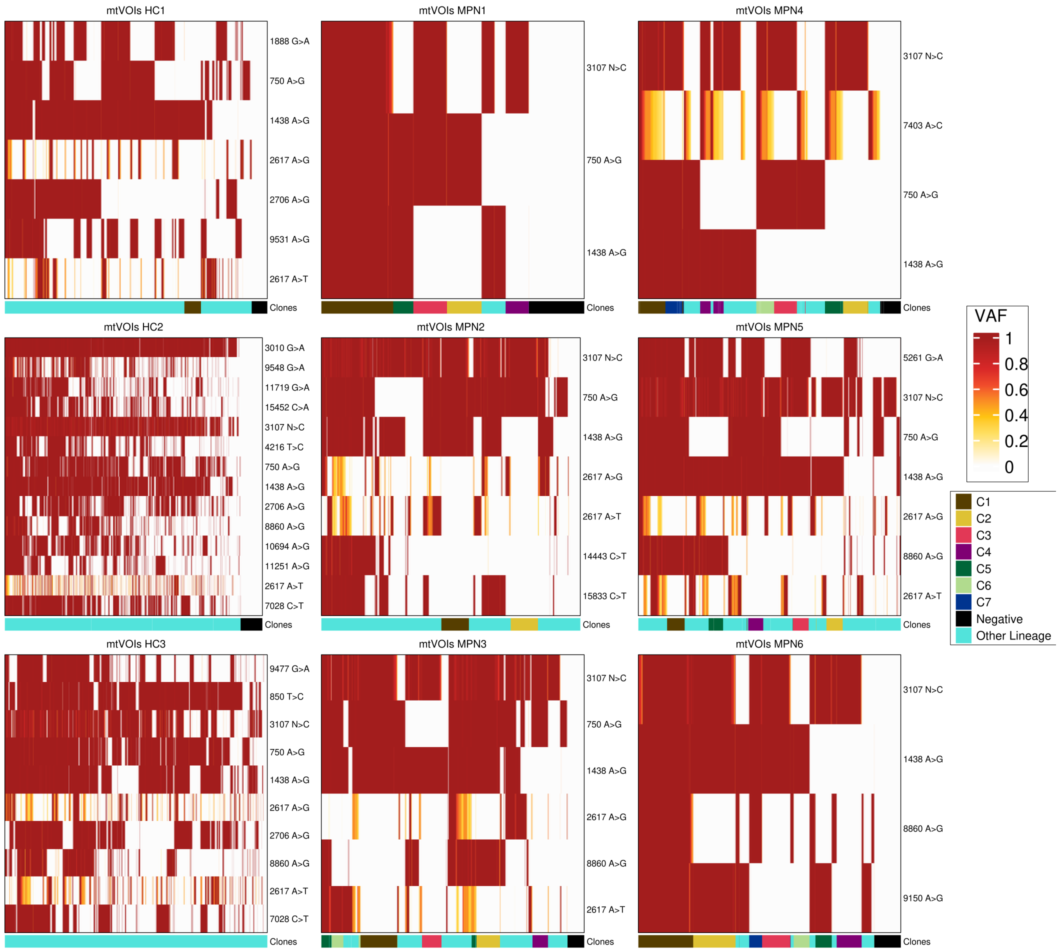


**Figure S6. mtVOIs per Sample.** Heat map showing the allele burden for mtVOIs selected for all samples. The bar below indicate identified clones. For convenience, only the top 7 largest clones and the negative clone have been colored, while other clones have grouped as Other Lineage.

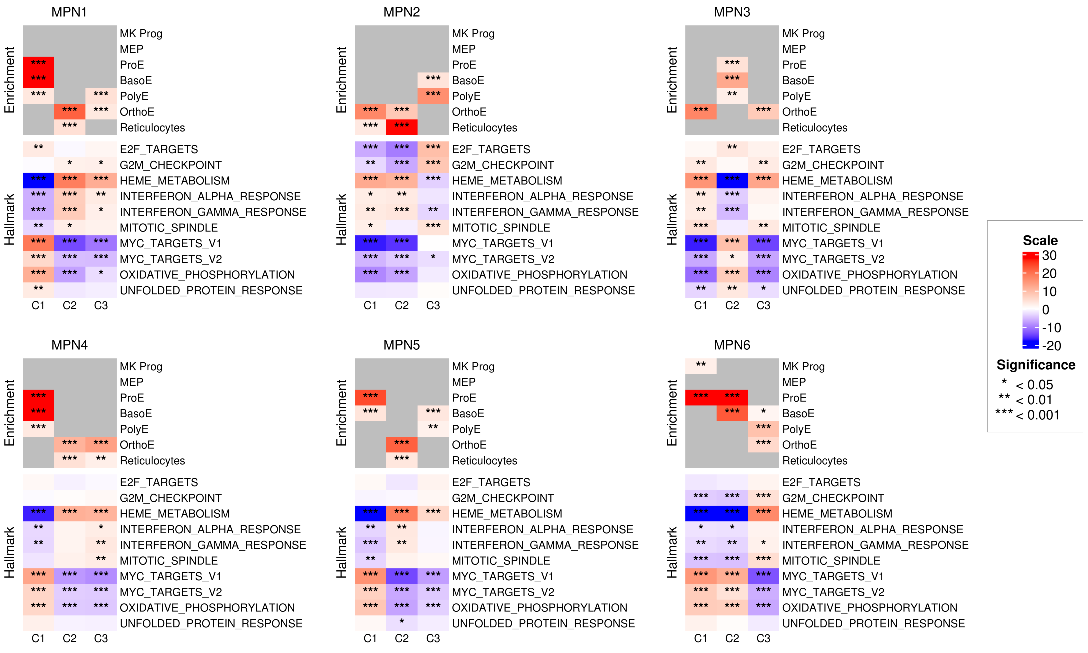


**Figure S7.** **Cell Type Enrichment and Hallmark GSEA for top 3 mtVar based clones for all MPN samples.**

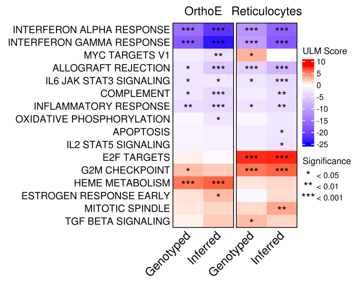


**Figure S8.** **Hallmark Pathways for *JAK2V617F* Positive and *JAK2*V617F inferred cells.**

**Table S1. Experimental Design Matrix.** The design matrix contains the information necessary to analyse the genotyping data. The individual column indicates to which patient/donor a sample belongs. The Sample column indicates the experimental condition of the measurement (treatment vs control). The Source column indicates which tools was used for the genotyping. The Type column indicates what type of sequencing data is used. The BAM, Molecule Info and Cell Barcodes columns indicate the location of the filtered alignment files for genotyping. The Results Path column is the output folder of the genotyping tools. Additional columns indicate the path of barcodes or molecule files.

| Individual | Sample | Source | Type | Results Path | BAM | Cell Barcodes | Molecule Info |
|---|---|---|---|---|---|---|---|
| MPN1 | MPN1_C | VarTrix | Amplicon | MPN1_C/ | MPN1_C_Filtered.bam | MPN1_C_barcodes.tsv | MPN1_C.h5 |
| MPN1 | MPN1_C | VarTrix | scRNAseq | MPN1_C/ | MPN1_C_Filtered.bam | MPN1_C_barcodes.tsv | MPN1_C.h5 |
| MPN1 | MPN1_C | MAEGATK | scRNAseq | MPN1_C/maegatk.rds | MPN1_C_Filtered.bam | MPN1_C_barcodes.tsv | MPN1_C.h5 |
| MPN1 | MPN1_T | VarTrix | Amplicon | MPN1_T/ | MPN1_T_Filtered.bam | MPN1_T_barcodes.tsv | MPN1_T.h5 |
| MPN1 | MPN1_T | VarTrix | scRNAseq | MPN1_T/ | MPN1_T_Filtered.bam | MPN1_T_barcodes.tsv | MPN1_T.h5 |
| MPN1 | MPN1_T | MAEGATK | scRNAseq | MPN1_T/maegatk.rds | MPN1_T_Filtered.bam | MPN1_T_barcodes.tsv | MPN1_T.h5 |

**Table S2. Number of cells genotyped for *JAK2*V617F per cell type.**

| CellType | Positive | Negative | NoCall |
|---|---|---|---|
| MK Progenitors | 2 | 0 | 69 |
| Proerythroblasts | 109 | 212 | 6,136 |
| MEP | 2 | 1 | 685 |
| Basophilic Erythroblasts | 83 | 118 | 5,506 |
| Polychromatic Erythroblasts | 144 | 144 | 8,334 |
| Orthochromatic Erythroblasts | 201 | 144 | 13,147 |
| Reticulocytes | 49 | 25 | 3,880 |
| Monocytes | 1 | 16 | 291 |
| Mast Cell Progenitors | 2 | 0 | 261 |
| Granulocytes | 6 | 28 | 775 |
| Macrophages | 2 | 10 | 131 |
| T Cells | 2 | 3 | 356 |
| B Cells | 0 | 0 | 74 |
